## Supplementary data 5 for "The Idiopathic Pulmonary Fibrosis-associated single nucleotide polymorphism rs35705950 is transcribed in a MUC5B Promoter Associated Long Non-Coding RNA (AC061979.1)"

### –Regions Under eXploration– Mining RNA-SEQ Data for Novel RNA Transcripts

**Ruxandra Neatu MSc**

Northumbria University
Department of Applied Sciences
Newcastle, UK
January 2020

### Summary

**Introduction** **2**

**1 RNA-SEQ pipeline** **3**

**2 Public RNA-SEQ Data Analysis in a Nutshell** **11**

**3 Example** **12**

**4 One More Thing** **15**

**References** **16**

### Introduction

This RNA-SEQ pipeline is designed for the rare transcript detection in the human genome by analysing publicly available data. The process is divided in three sections:

*a)* **Obtaining** the RNA-SEQ studies and the corresponding SRP accession number.

*b)* **Processing** the data: Galaxy Project or local cluster[1].

*c)* **Collapsing** the studies.

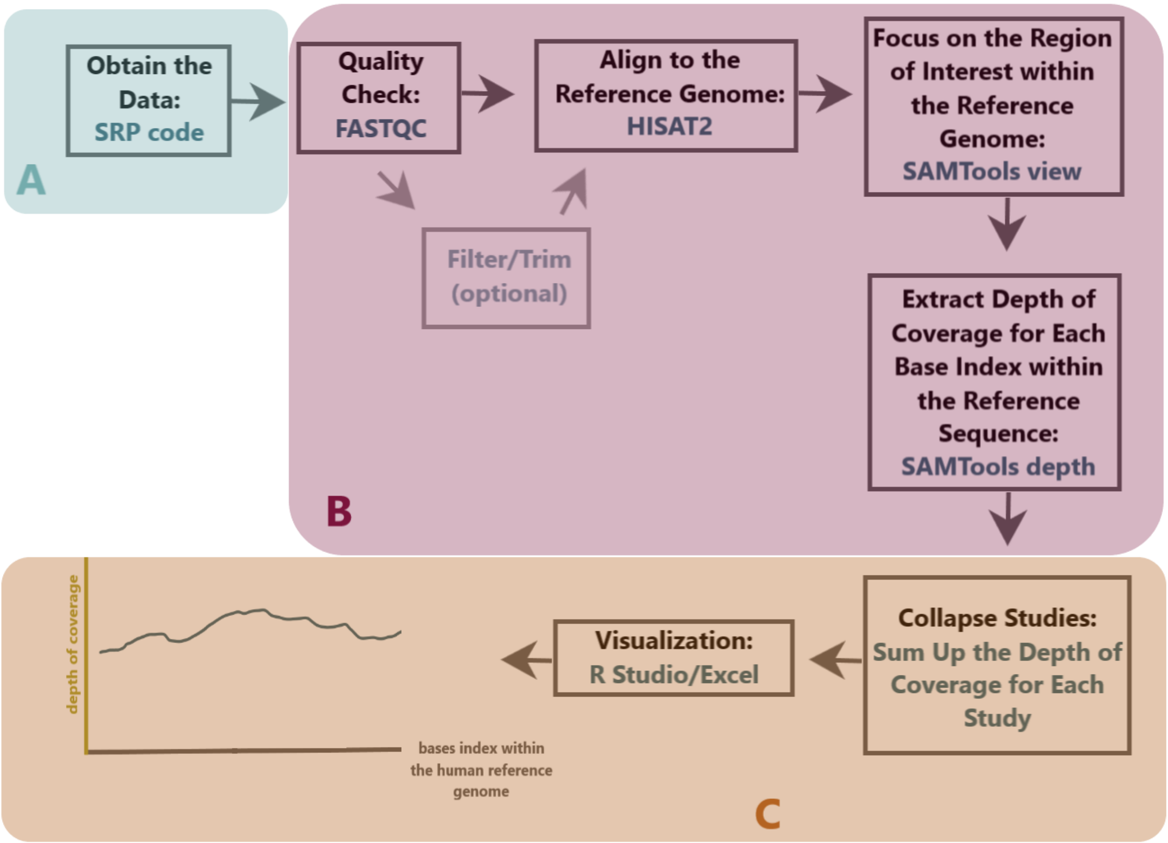

Figure 1: RNA-SEQ pipeline overview

The method can be applied globally in the DNA. However, in this case,the target of the pipeline will be the intergenic region between MUC5AC-MUC5B protein-coding genes (1,201,139 - 1,223,065 on chromosome 11)[2]. The method was developed for the detection of RNA transcripts that align on the rs35705950 polymorphism (locus:1,219,991)[2] in the human genome. Figure 2 is a simplified diagram of the NCBI gene database (site here[3]).

[1]Only free resources will be used here: Section B will be managed entirely by Galaxy Project.

[2]Genomic Sequence: NC_000011.10 Chromosome 11 Reference GRCh38.p13, the most recent one to date

[3]Links are available under the word *here* throughout the document

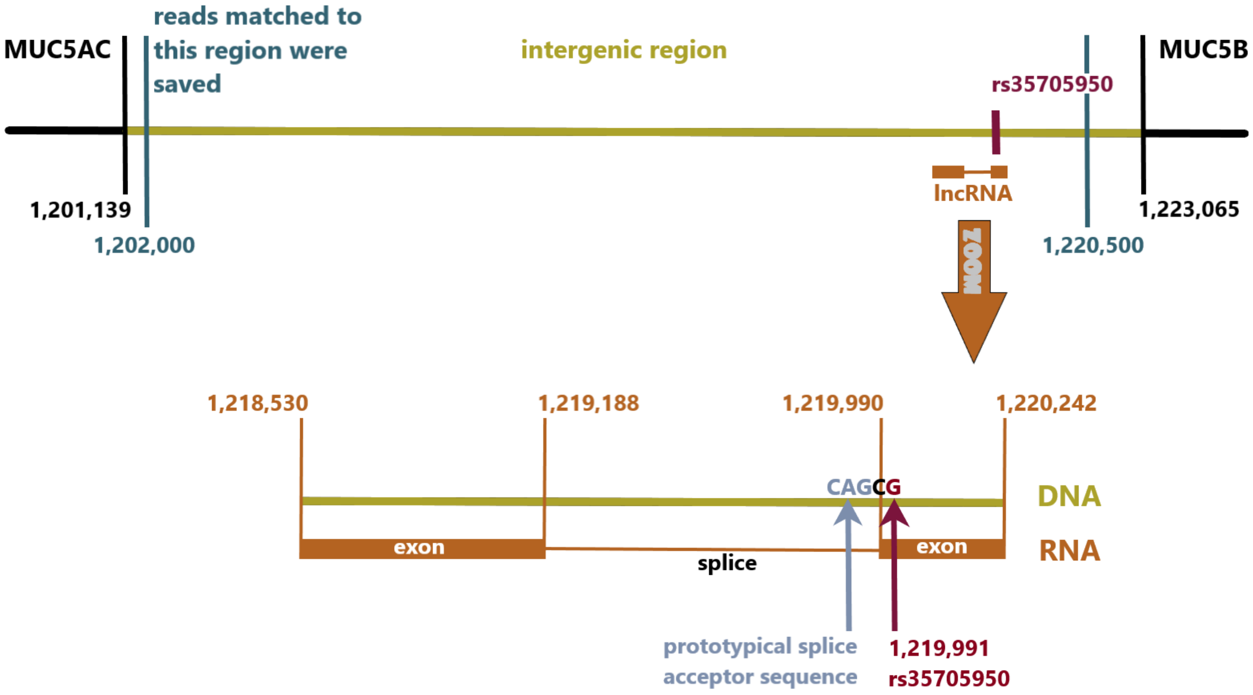

Figure 2: MUC5B-MUC5AC intergenic region with focus on the rs35705950 SNP and the lncRNA. Region 1,202,000 - 1,220,500 (blue) represents the Focused BAMs (see 1.2.2).

### RNA-SEQ pipeline

#### 1.1 Section A: Obtaining Data

- **STUDIES**
  - Click here (NCBI) → Database: PubMed → Search (keywords): RNA-SEQ A549/BEAS2B/lung epithelial cells[4].
  - Select study.
- **SRA accession number**
  - Identify the SRP accession number (usually in the "Materials and Methods").
  - Sometimes only a GEO accession number is provided. In this case refer to chapter 3: Example.
- **SRA Run Selector**
  - Click here (NCBI SRA Run Selector).

[4] Our research centres hereditary lung diseases. The keywords depend on the aim of the meta-analysis.

– Insert the SRA accesion number.

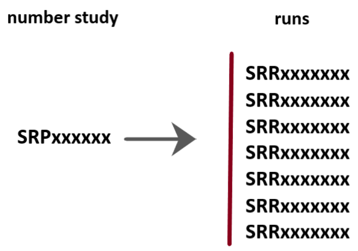

Figure 3: A SRA accesion number (SRPxxxxxx) includes the runs (SRRxxxxxx) within the study

– Download the SRR Accesion list (tabular file .TXT). It contains the individual runs (see *Public RNA-SEQ Data Analysis in a Nutshell* at the end of the document).

#### 1.2 Section B: Galaxy Project

##### 1.2.1 Uploading the Data in Galaxy

- **Create an account.** You must have an account in order to be able to use it: link here.
- **Upload the .TXT file** (SRR Accession list) downloaded in SECTION A.
  - Galaxy path: Get Data → Upload file
- **Upload the actual runs from the .TXT file** into Galaxy for further manipulation.
  - Galaxy path: Get Data → Download and Extract Reads in FASTA/Q
    * Input type: List of SRA accesion, one per line → Select the SRR Accesion list[5].

##### 1.2.2 Processing the Data: HISAT2 and SAMTools

- **FASTQC.**[6] After the data is uploaded to the Galaxy server, it is good practice to use FASTQC for checking the raw reads (especially for the publicly available ones).
  - Galaxy path: Genomic File Manipulation → FASTQ Quality Control → FastQC Read Quality report.
- **HISAT2.** Aligning the reads to the human reference genome.

[5]Important: Either Galaxy or a local cluster, the storage space is crucial (max. 200GB for Galaxy). If there are too many runs in the list (add up the space needed for the output of the analysis), the storage space might be exceeded and the job will end with an error. In this case, the best practice is to work in batches. The resulting files can be merged as desired (see 1.2.2 SAMTools merge)

[6]For the tools used in Galaxy: only the parameters that will be changed are mentioned. Otherwise assume default settings (set up by Galaxy).

– Galaxy path: Genomic analysis → RNA-seq → HISAT2
  * Use a built-in genome: Human (*Homo sapiens)*) (b38):hg38
  * Library type: Single-end or Paired-end (Default values)
  * Strandness: Unstranded, Forward (FR), Reverse(RF)
– For *Library type* and *Strandness*: it depends on the RNA-SEQ library preparation (Strandness) and the sequencing (Library Type). The information is available in the *Materials and Methods* of each study (see chapter 3: Example).

• **SAMTools view.**[7]

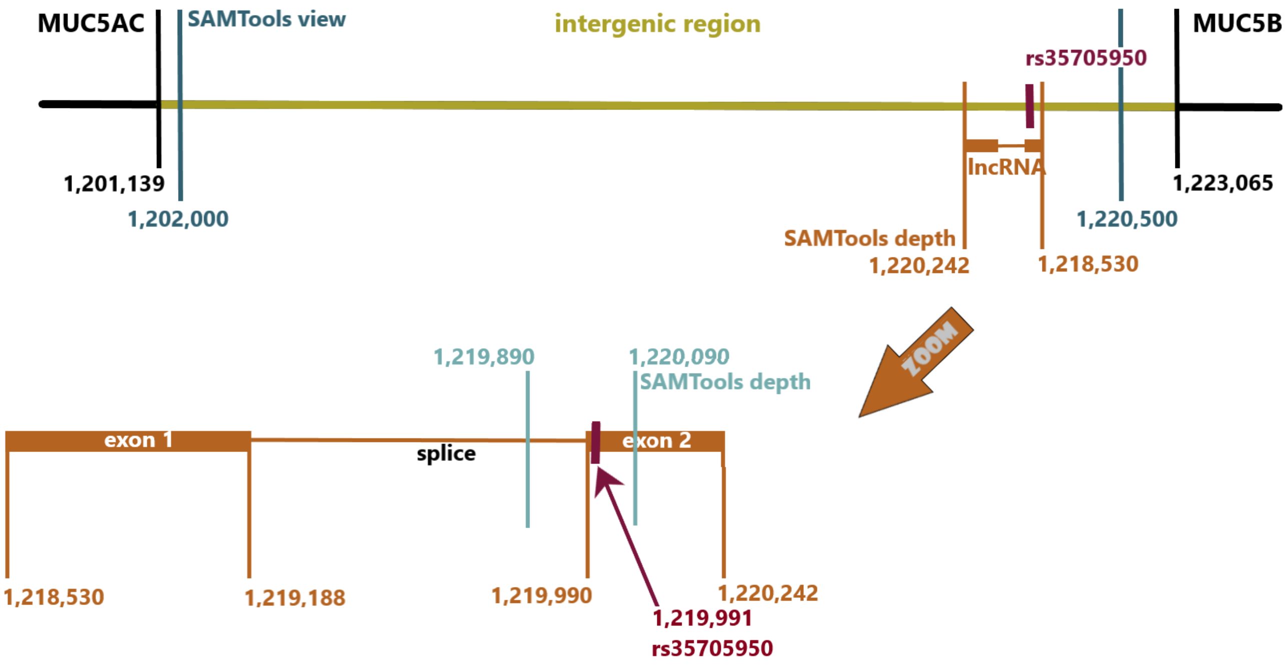

Figure 4: SAMTools view and SAMTools depth will work on the marked regions. These positions are specific to the reference sequence (in this case chromosome 11, *Homo sapiens*[8]).

– Galaxy path: Genomic file manipulation → SAM/BAM → Samtools view
  * Input: the files resulted from HISAT2. There are BAM (Binary Alignment Map) format
  * Output type: BAM (-b)
  * Filter alignment: yes
  * Filter by region: manually specify regions: chr11:1,202,000-1,220,500 or chr11:1,220,242-1,218,530
– As in figure 5 (yellow), the output of SAMTools view is considerably smaller thus easier to store and manipulate. This is because only reads that matched to the 20,500 nt are stored.

[7]This step is required for storing the data and further manipulating it on a personal computer(see figure 5 and 1.2.3)
[8]Genomic Sequence: NC_000011.10 Chromosome 11 Reference GRCh38.p13, the most recent one to date

- **SAMTools merge.**
  - Galaxy path: Genomic file manipulation → SAM/BAM → Samtools merge
  - A study can have hundreds of runs. They can be put together in a single file. At this step, the BAM files (before or after merging) can be visualised in Integrative Genomic Viewer (IGV).
  - Note: SAMTools merge makes the data easier to manipulate since it concatenates all of the reads in a single file. This is instead of running the same process on each individual run (some studies can have hundreds). However, it is important to have the BAM for each run (see 1.2.4. BAM Files Storage) (e.g.: a mutation (visualised in IGV) can be in some samples but not in others).

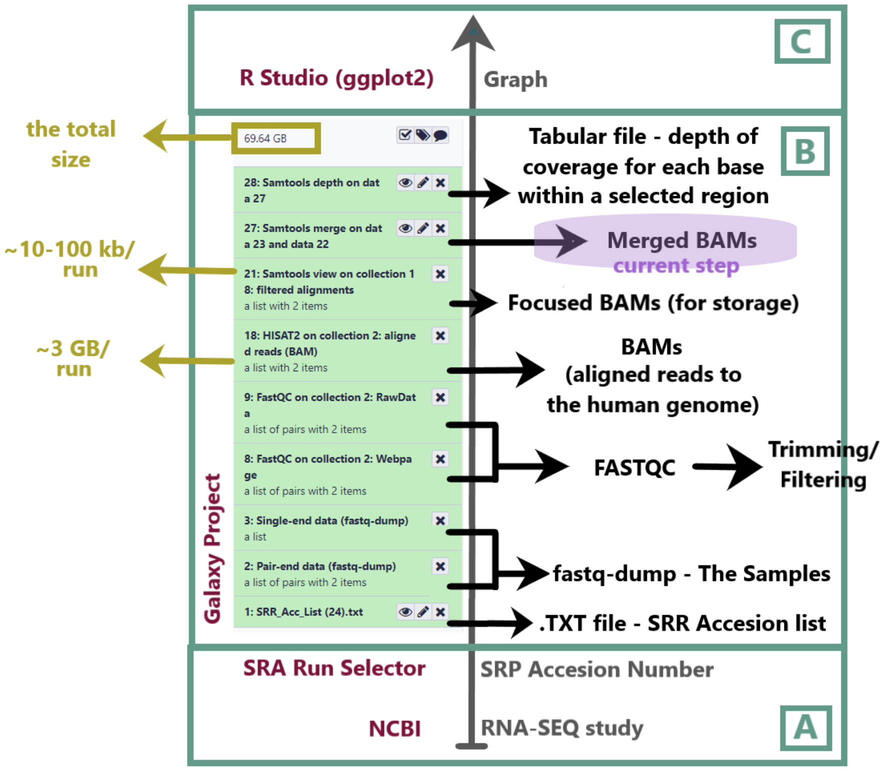

Figure 5: Galaxy History Overview. By the end the history should look like this. The purple highlight represents the current step.

In Galaxy, it is possible to go from HISAT2 output (BAMs) to the SAMTools depth (to extract depth of coverage for each position/base). They are treated the same as the Focused BAMs. The SAMTools depth will only take into consideration the DNA region that is specified[9]. However, because of the size (see figure 5, yellow), it might be convenient to save only the reads that match the DNA region that serves a specific research purpose. From then on, the Focused BAMs can be manipulated on any computer with SAMTools/BAMTools installed.

[9]Once you have the Focused BAMs, SAMTools depth will only output zero values in the regions exceeding the ones specified in SAMTools view.

##### 1.2.3 Extract the Depth of Coverage

- **SAMTools depth.** This step can be achieved in one of two ways: Galaxy Project or any computer with SAMTools/BAMTools installed.
  - **Option 1**[10]**.** Galaxy path: Genomic file manipulation → SAM/BAM → Samtools depth
    - Output all positions: -aa
    - Filter by region: chr11:1,219,890-1,220,090 or the exact length of the lncRNA (figure 4) chr11:1,218,530-1,220,242
    - Input: BAMs or the focused BAMs
  - **Option 2**[11]**.** SAMTools command line[12]:

```
samtools depth -aa -r chr11:1,219,890-1,220,090 input.bam > output.txt
```

  - Both methods will create a tabular file with 3 columns: the name of the reference sequence, the base index within the reference, the depth of coverage for that base.

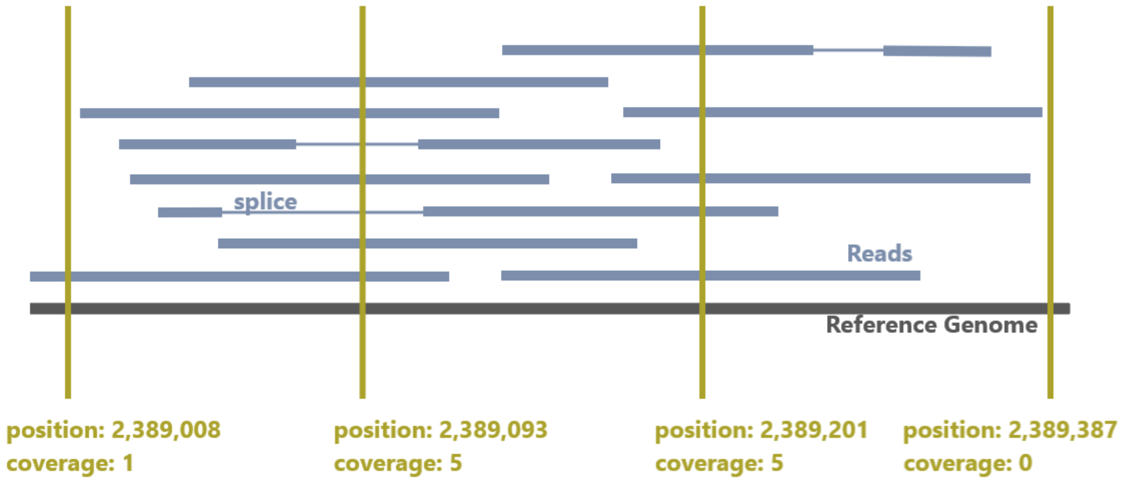

Figure 6: Illustration SAMTools depth. Splices are ignored by default.

##### 1.2.4 BAM Files Storage

The *Focused BAMs* (resulted from SAMTools view and SAMTools merge) files are downloaded and stored (if desired) as explained below:

[10]SAMTools depth can work directly on the HISAT2 output without SAMTools view.
[11]It requires the SAMTools view step and for the focused BAMs to be downloaded from the Galaxy Cluster. In case of a large meta-analysis, 1.2.4 explains their storage.
[12]For more options refer to the manpage here

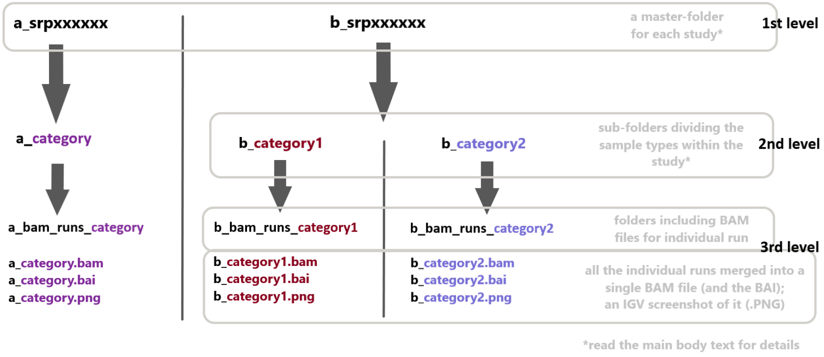

Figure 7: Processed RNA-SEQ Data Summary. Legend: a, b – assigned numbers corresponding to each study, consistent throughout the process for an easier manipulation of the data; srpxxxxxx – SRA accession number for each study

- **The datasets are divided firstly by study.** For each study there is an assigned number (consistent throughout documents) and an SRA accession number which represents the name of the master-folders (e.g.: 04_srp060719). Folders that have the same SRA accession number but a different assigned number is an indication that different parameters have been used on the same data.
- **Within the master-folder** of each study:
  - Samples are divided in different categories (see b_srpxxx xxx in Figure 1) or not (see a_srpxxxxxx in Figure 1). The default criteria for sorting is by sample type (A459, BEAS2B, NHBE etc.) but can also be by the state of the patient disease (non-CF, CF) or just by patient (donor A, donor B).
  - Each scientific article is going to have a different sorting system due to their great diversity. For each category is a different sub-folder (see b_category1 and b_category2 in the figure 7).
  - **Within the sub-folder:**
    - A folder with individual BAMs for each run.
    - And additional files: a single BAM files with all the individual runs merged and the corresponding BAI; an IGV screenshot of it (PNG file).

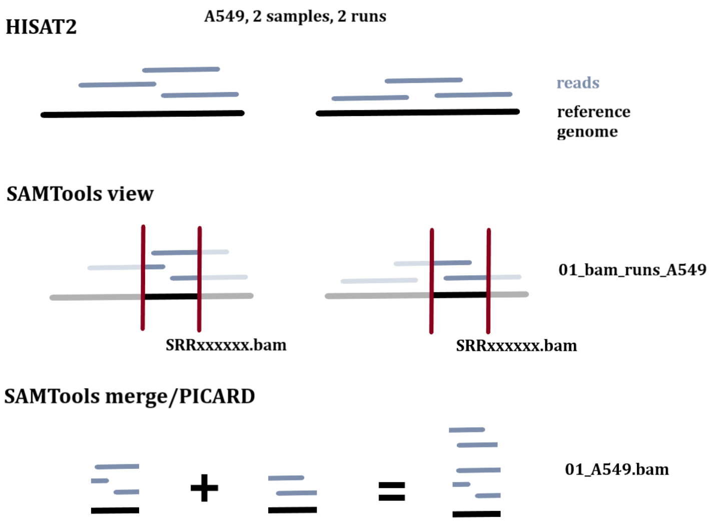

Figure 8: Example of the files content within the 3rd level (assuming only two runs from A549 cells (for the clarity)).

#### 1.3 Section C: Collapse Studies

- **R Studio(ggplot2)** - Visualisation

Final output from the previous step: .TXT file with 3 columns.

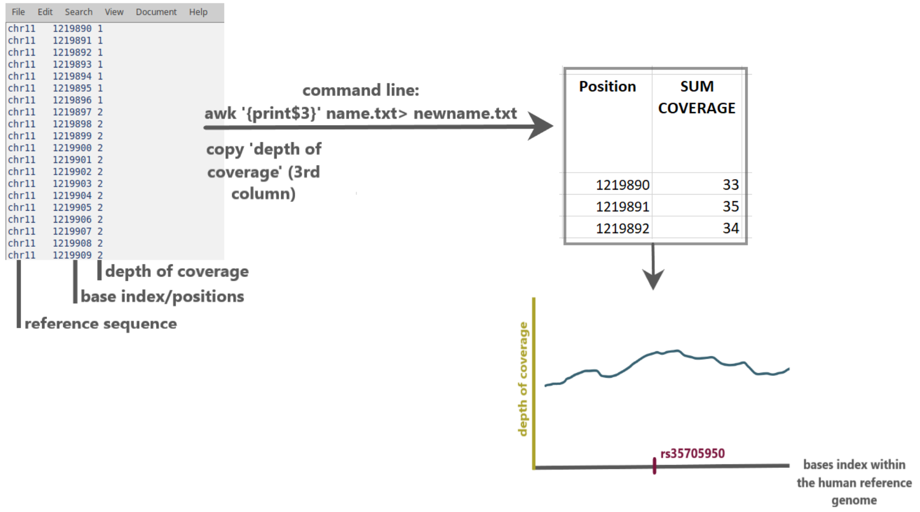

Figure 9: Final step of the analysis includes summing up the depth of coverage for all the studies.

As in figure 9, the depth of coverage (third column) is extracted into a new tabular file (for the convenience) using the awk command then paste it in excel. The third column (depth of coverage) is summed up from the samples across all of the studies. R Studio, package ggplot2, was used to generate the graph. Code available upon request. Alternatively Excel can offer a fast preview of the data.

Once extracted, the depth of coverage for each position can be summed up and compared in many different ways (see 3. Example). For example, in this meta-analysis, the depth of coverage from epithelial and basal cells within the same study have been compared. Similarly depth of coverage from the BEAS2B (bronchial normal cells) and A549 (bronchial cancerous cells) cell lines across different studies can be compared. This depends on the aims of the research.

### Public RNA-SEQ Data Analysis in a Nutshell

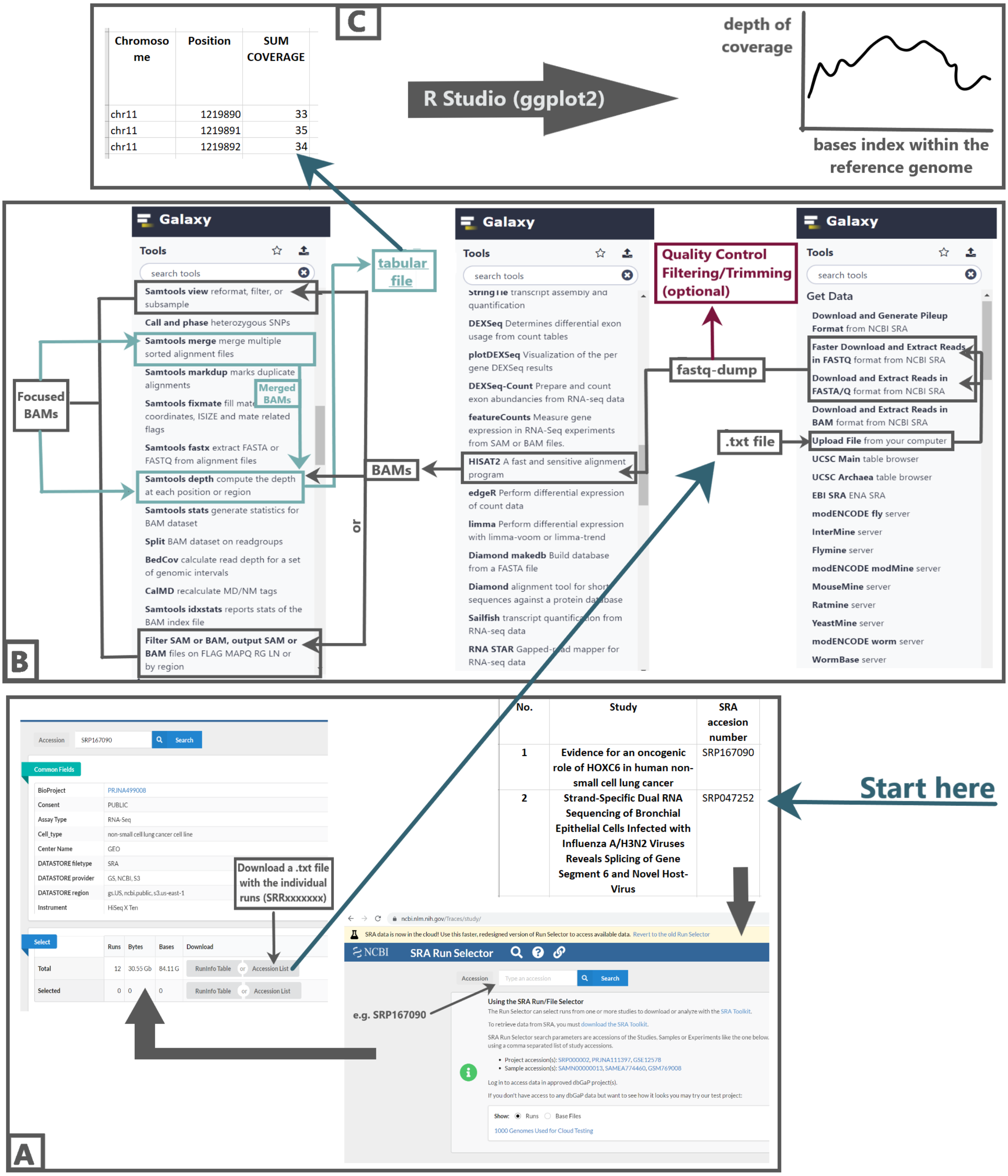

### Example

1. Study: Zhang *et al.* 2018[1]

2. GEO Accesion number: GSE86064 (Omnibus site here) – found in the Methods → Clinical samples and RNA sequencing

3. In the bottom of the Omnibus site: SRA Run selector link. This will redirect to here. Otherwise, on the same page the BioProject and SRA accesion number (for this study: SRP082973) are also available.

4. From SRA Run Selector, select the runs on the basal cells. Download the .TXT file. This will only contain the SRR accesion numbers for those runs. It is the same for the epithelial cells.

5. **Galaxy Project:** A .TXT files that conains only runs from one type of cells will be uploaded in Galaxy (Section B: 1.2.1). Galaxy has a storage limit. If the files that follow to be uploaded are exceeding this limit, the job will end with an error. In this case divide the SRR accesion numbers in more .TXT files and process them separately. They can be merged together at the end (see 1.2.2 SAMTools Merge).

6. **Strandness[13]: Unstranded, Forward, Reverse.** The strandness depends on the cDNA Library Preparation.

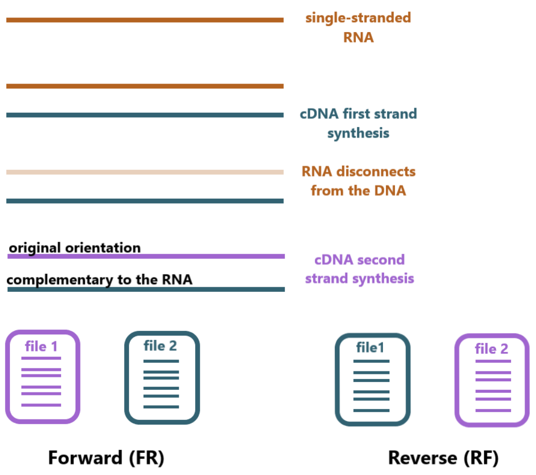

Figure 10: Simplified diagram of an RNA-SEQ library preparation. Forward(FR) and Reverse(RF) are specific to HISAT2 (TopHat, for example, has the option first-strand and second-strand but it is the same concept). When reads corresponding to the initial orientation of RNA are stored in file 1 and the reads from the complementary strand in file 2 → Forward (FR). When reads corresponding to the complementary strand are stored in file 1 and the reads from the initial strand are stored in file 2 → Reverse (RF). Unstranded → the reads can originate from both sense and antisense strands thus is less specific, however, HISAT2 manages it well.

[13]The explanation applies only to paired-end data. The same rule applies to single-end with the exception that file 2 does not exist thus only one of the two strands will be sequenced.

This information is extracted from the sequencing protocol (not all studies have this available[14]). In this case, the sequecing protocol is available in the Supplementary information of the study: TruSeq RNA Sample Prep Kit (Illumina San Diego, CA) which is non-stranded (choose option *Unstranded* when running HISAT2 for this dataset.) See documentation here.

7. **Library type: single-end or paired-end?** The library type depends on the cDNA sequencing. This information is availabe in Methods → Clinical samples and RNA sequencing. Additionally Galaxy will automatically separate single-end and paired-end runs (fastq-dump).

   Note: If more than one run is uploaded at once, choose Paired-end Dataset Collection.

8. After processing the runs originated from the epithelial cells and from basal cells, they should be merged by type. The depth of coverage from epithelial cells can be compared with the depth of coverage from the basal cells. This can be visualised as explained in 1.3. Section C.

   A table with the SRR accesion numbers (LA - large airway; SA - small airway) for the samples provided by Zhang *et al.* 2018 study is available below.

[14] In this case, *Unstranded* option will suffice

Table 1: Zhang *et al.* 2018 Samples

| | Source | | | |
|---|---|---|---|---|
| | LA epithelial cells | LA basal cells | SA epithelial cells | SA basal cells |
| SRR acc. | SRR4064971 | SRR4064981 | SRR4065001 | SRR4065024 |
| | SRR4064972 | SRR4064982 | SRR4065002 | SRR4065025 |
| | SRR4064973 | SRR4064983 | SRR4065003 | SRR4065026 |
| | SRR4064974 | SRR4064984 | SRR4065004 | |
| | SRR4064975 | SRR4064985 | SRR4065005 | |
| | SRR4064976 | SRR4064986 | SRR4065006 | |
| | SRR4064977 | SRR4064987 | SRR4065007 | |
| | SRR4064978 | SRR4064988 | SRR4065008 | |
| | SRR4064979 | SRR4064989 | SRR4065009 | |
| | SRR4064980 | SRR4064990 | SRR4065010 | |
| | | SRR4064991 | SRR4065011 | |
| | | SRR4064992 | SRR4065012 | |
| | | SRR4064993 | SRR4065013 | |
| | | SRR4064994 | SRR4065014 | |
| | | SRR4064995 | SRR4065015 | |
| | | SRR4064996 | SRR4065016 | |
| | | SRR4064997 | SRR4065017 | |
| | | SRR4064998 | SRR4065018 | |
| | | SRR4064999 | SRR4065019 | |
| | | SRR4065000 | SRR4065020 | |
| | | SRR5445518 | SRR4065021 | |
| | | SRR5445519 | SRR4065023 | |
| | | SRR5445520 | | |
| | | SRR5445521 | | |
| | | SRR5445522 | | |
| | | SRR5445523 | | |
| | | SRR5445524 | | |
| | | SRR5445525 | | |
| | | SRR5445526 | | |
| | | SRR5643321 | | |
| | | SRR5643322 | | |
| | | SRR5643323 | | |
| | | SRR5643324 | | |
| | | SRR5643325 | | |
| | | SRR5643326 | | |
| | | SRR5643327 | | |
| | | SRR5643328 | | |
| | | SRR5643329 | | |
| | | SRR5643330 | | |
| | | SRR5643331 | | |
| | | SRR5643332 | | |
| | | SRR5643333 | | |

### One More Thing

Document available at github.com/NeatuR/RUX-RNA-SEQ. This is part of a larger project that aims to explain the biological role of promoter associated long non-coding RNAs.

✉
